# The *Shigella* type III effector protein OspB is a cysteine protease

**DOI:** 10.1101/2021.09.01.458261

**Authors:** Thomas E. Wood, Kathleen A. Westervelt, Jessica M. Yoon, Heather D. Eshleman, Roie Levy, Henry Burnes, Daniel J. Slade, Cammie F. Lesser, Marcia B. Goldberg

## Abstract

The type III secretion system is required for virulence of many pathogenic bacteria. Bacterial effector proteins delivered into target host cells by this system modulate host signaling pathways and processes in a manner that promotes infection. Here, we define the activity of the effector protein OspB of the human pathogen *Shigella* spp., the etiological agent of shigellosis and dysenteric disease. Using the yeast *Saccharomyces cerevisiae* as a model organism, we show that OspB sensitizes cells to inhibition of TORC1, the central regulator of growth and metabolism. *In silico* analyses reveal that OspB bears structural homology to bacterial cysteine proteases, and we define a conserved cysteine-histidine catalytic dyad required for OspB function. Using yeast genetic screens, we identify a crucial role for the arginine N-degron pathway in the growth inhibition phenotype and show that inositol hexakisphosphate is an OspB cofactor. We find that a yeast substrate for OspB is the TORC1 component Tco89p, proteolytic cleavage of which generates a C-terminal fragment that is targeted for degradation *via* the arginine N-degron pathway; processing and degradation of Tco89p is required for the OspB phenotype. In all, we demonstrate that the *Shigella* T3SS effector OspB is a cysteine protease and decipher its interplay with eukaryotic cell processes.

**Importance:** *Shigella* spp. are important human pathogens and one of the leading causes of diarrheal mortality worldwide, especially in children. Virulence depends on the *Shigella* type III secretion system (T3SS), and definition of the roles of the bacterial effector proteins secreted by the T3SS is key to understanding *Shigella* pathogenesis. The effector protein OspB has been shown to contribute to a range of phenotypes during infection, yet the mechanism of action is unknown. Here, we show that OspB possesses cysteine protease activity in both yeast and mammalian cells, and that enzymatic activity of OspB depends on a conserved cysteine-histidine catalytic dyad. We determine how its protease activity sensitizes cells to TORC1 inhibition in yeast, finding that OspB cleaves a component of yeast TORC1, and that the degradation of the C-terminal cleavage product is responsible for OspB mediated hypersensitivity to TORC1 inhibitors. Thus, OspB is a cysteine protease that depends on a conserved cysteine-histidine catalytic dyad.

## Introduction

Cellular processes are largely controlled by both the availability of nutrients and the ability to respond to these environmental cues. Consequently, homeostatic control of metabolism is crucial to function, growth and ultimately viability. The balance between anabolic and catabolic processes in eukaryotes is controlled by the target of rapamycin (TOR) complex, a large multi-subunit hub integrating sensory inputs to regulate cellular metabolism. In nutrient-replete conditions, amino acid and glucose availability sustains the kinase activity of the TORC1 complex, promoting translation, gene expression and protein stability. In contrast, cellular stresses and starvation inhibit TORC1-dependent growth and stimulate proteolytic mechanisms such as autophagy to maintain amino acid pools (González & Hall, 2017).

Infection perturbs cellular homeostasis (Eisenreich et al., 2013). Pathogens that invade host cells disrupt cellular processes in ways that promote the survival and replication of the infectious agent. *Shigella* spp. are the etiological agent of bacillary dysentery and a leading contributor to diarrheal mortality (Khalil et al., 2018). This pathogen invades the intestinal epithelium, establishing a replicative niche within colonic epithelial cells and triggering an acute inflammatory immune response (Carayol & Tran Van Nhieu, 2013). The type III secretion system (T3SS) is required for *S. flexneri* infection, facilitating invasion and bacterial replication through the delivery into host cells of effector proteins that subvert cellular signaling pathways. Effector proteins also promote the spread of intracellular *Shigella* between cells, whereby it disseminates throughout the intestinal epithelium (Agaisse, 2016).

*Shigella* T3SS effector proteins display a myriad of well-characterized enzymatic activities including phosphatase, acyltransferase, ubiquitin ligase and protease functions, as well as catalyzing other more unconventional biochemical modifications (Burnaevskiy et al., 2013; H. Li et al., 2007; Z. Li et al., 2021; Liu et al., 2018; Niebuhr et al., 2002; Rohde et al., 2007; Zhang et al., 2012). The effector OspB has been described by our group and others as manipulating mTORC1 signaling, dampening the innate immune response *via* MAP kinase and NF-κB signaling, and modulating cytokine release (Ambrosi et al., 2015; Fukazawa et al., 2008; Zurawski et al., 2009). However, the precise mode of action of this effector protein is unknown.

In this study, we determine that the *S. flexneri* T3SS effector OspB is a cysteine protease. *In silico* analysis indicated structural homology to bacterial cysteine proteases, permitting identification of putative catalytic residues. Using a yeast model to study the impact of OspB activity on a eukaryotic host, we determine host factors required for OspB-dependent hypersensitivity to TORC1 inhibition, including inositol phosphate biosynthesis, TORC1 signaling, and protein degradation pathways. We find that the OspB-dependent hypersensitivity phenotype is due to the cleavage of the TORC1 component Tco89p, in a manner that requires both the conserved catalytic dyad in OspB and the secondary messenger molecule inositol hexakisphosphate. Finally, we demonstrate that the C-terminal product of Tco89p cleavage enters the arginine N-degron pathway for destruction by the proteasome, and that its degradation is required for OspB-dependent growth inhibition of yeast.

## Results

### OspB exhibits structural homology to cysteine proteases

To gain insight into the potential mechanism of action of OspB, we performed *in silico* analyses of its amino acid sequence. This analysis revealed that OspB shares 27-30% sequence identity with the cysteine protease domains (CPD) of the large clostridial cytotoxins TcdA and TcdB of *Clostridioides difficile* and the multifunctional autoprocessing repeats-in-toxin (MARTX) RtxA toxins of *Vibrio cholerae* and *V. vulnificus* (**Fig. 1A**). TcdA, TcdB and RtxA are modular toxins that upon host cell endocytosis undergo autoproteolysis, which releases toxin domains that subvert cellular processes by inducing actin depolymerization and altering GTPase signaling (Fullner & Mekalanos, 2000; Just et al., 1995). In contrast to these large cytotoxins, OspB is small (288 amino acids; 32 kD), and we found no evidence for OspB autoprocessing in cells (**Fig. S1** in the supplemental material).

**FIGURE 1:**
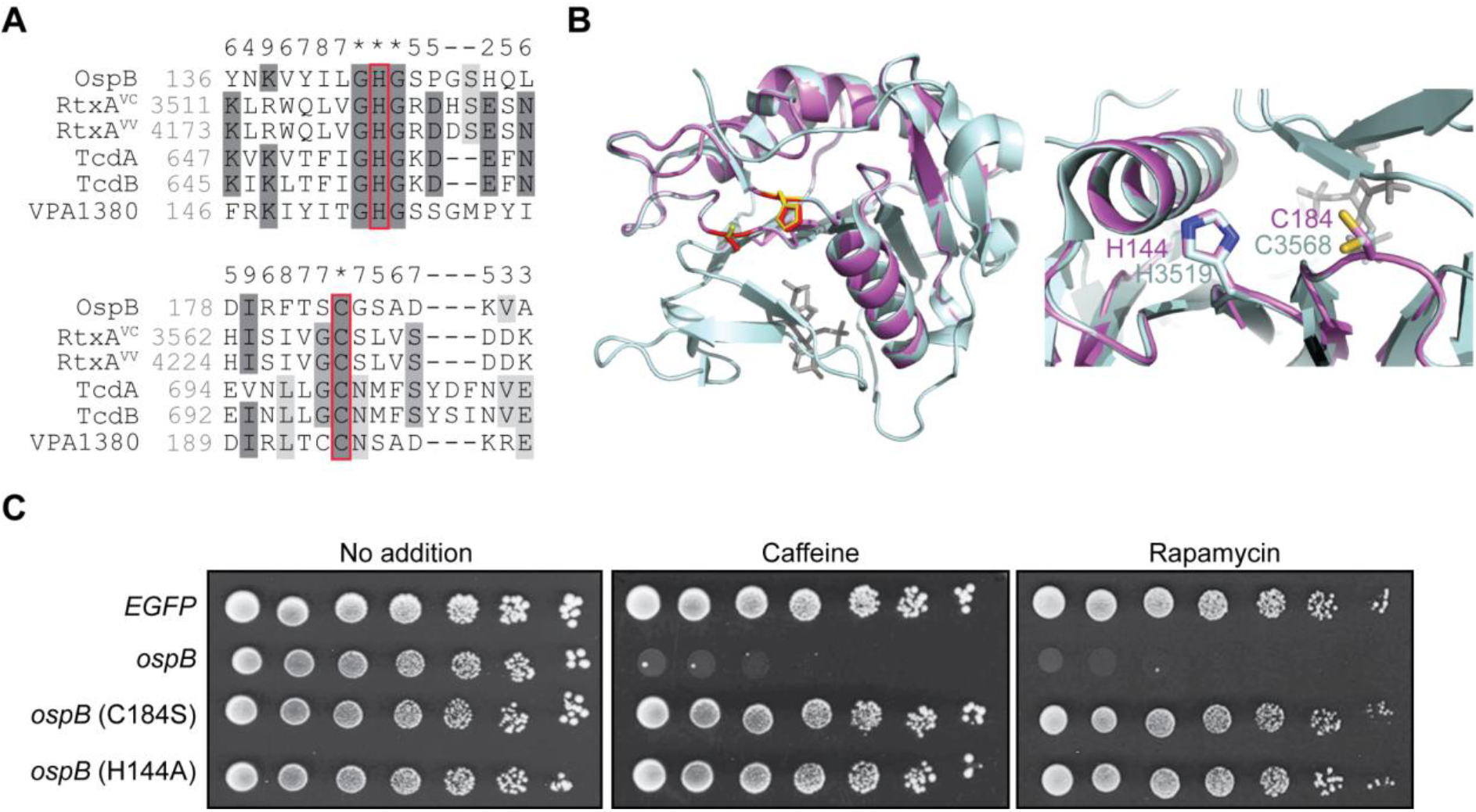
OspB possesses a predicted cysteine protease catalytic dyad. (A) Multiple sequence alignment of OspB with the catalytic residues of the cysteine protease domains of RtxA from *V. cholerae* (RtxA^VC^) and *V. vulnificus* (RtxA^VV^), *C. difficile* TcdA and TcdB, and the OspB ortholog VPA1380 from *V. parahaemolyticus*. Red boxes indicate catalytic residues of the cysteine protease domains and the aligned putative catalytic residues of OspB. Darkness of gray shading reflects the conservation of individual residues, and the numbers above the alignment score the conservation at each position. Asterisks denote full conservation among the aligned sequences. (B) Cartoon depiction of a tertiary structure model of OspB (violet) on the CPD of RtxA^VC^ (*left panel*; PDB: 3EEB) (pale cyan). In the left panel, the catalytic residues of the cysteine protease domain are denoted by yellow sticks, with the putative catalytic residues of OspB shown as red sticks. The inositol hexakisphosphate cofactor in the RtxA^VC^ cysteine protease domain structure is shown in dark gray. In the right panel, an enlarged and rotated view shows the active site, highlighting the superposition of the putative OspB catalytic residues with those of the cysteine protease domain, labelled according to the color of the cartoon. (C) Growth of yeast strains expressing *ospB* constructs or an *EGFP* control. Serial dilutions were spotted on media either without additives or supplemented with the TORC1 inhibitors caffeine or rapamycin (*n* = 3).

The cysteine and histidine residues required for the proteolytic activity of the CPDs are conserved in OspB and the orthologous T3SS effector protein of *V. parahaemolyticus* VPA1380 (Calder et al., 2014; Egerer et al., 2007; Sheahan et al., 2007) (**Fig. 1A** and **Fig. S2A**). Indeed, the tertiary structure of OspB can be modelled on the CPDs of RtxA and TcdA with 96% and 62% confidence, respectively, with conservation of the positions of their catalytic residues with C184 and H144 of OspB (**Fig. 1B** and **Fig. S2B**). The alignment of OspB with the CPD structures suggested that OspB residues C184 and H144, may be required for OspB activity. A quantitative assay in yeast strains expressing *S. flexneri* effector proteins previously demonstrated that OspB causes growth inhibition of yeast in the presence of the cellular stressor caffeine (Slagowski et al., 2008). We utilized this assay to probe the role of the putative catalytic residues in OspB activity. Whereas expression of wild type OspB elicits a drastic growth defect in the presence of caffeine, mutation of either C184 or H144 completely abrogated toxicity (**Fig. 1C**). These data indicate that OspB inhibition of yeast growth depends on the predicted catalytic dyad of C184 and H144, bolstering our predictions for the tertiary structure of OspB as a structural homolog of the cysteine protease domains of several modular bacterial toxins.

Mutagenesis studies of TcdA showed that in addition to C700 and H655, D589 is required for autoprocessing through proton abstraction from the histidine in the active site (Pruitt et al., 2009). In RtxA, mutagenesis of the equivalent aspartic acid residue D3469 alone does not impact proteolytic activity; however concerted deletion of E3467 results in partial loss of autocleavage (Prochazkova & Satchell, 2008). In OspB an aspartic acid residue (D108) is predicted to be present at the equivalent position of D589^TcdA^, adjacent to two additional aspartic acid residues D109 and D110 (**Fig. S2A** and **S2B**). These were therefore candidates for involvement in catalysis. Alanine substitution of none of these aspartic acid residues rescued yeast growth (**Fig. S2C**). In addition, a D108A/D110A double mutant had no effect on OspB activity. These results indicate that the function of OspB requires both a cysteine and a histidine residue, similar to the cysteine-histidine catalytic dyad of the CPD of RtxA.

Among the effects of caffeine on cellular processes, inhibition of TORC1 is described as an important mode of action for this compound in yeast (Reinke et al., 2006). To determine whether the effect of caffeine on the OspB phenotype is specific to TORC1, we replaced caffeine in the media with rapamycin, which unlike caffeine is a specific inhibitor of TORC1. As with caffeine, the presence of rapamycin sensitized yeast to growth inhibition by OspB in a manner that depended on residues C184 and H144 (**Fig. 1C**). These data indicate that the impact of OspB on yeast growth depends on inhibition of TORC1, consistent with our previous findings that OspB potentiates rapamycin inhibition of growth in mammalian cells (Lu et al., 2015). The presence of similar OspB-dependent phenotypes in both yeast and mammalian cells with respect to sensitization to TOR inhibition demonstrates that yeast present a reasonable model for investigating the mechanism of OspB activity.

### Inositol hexakisphosphate is required for OspB activity

With the goal of identifying host factors required for OspB activity, we screened a *S. cerevisiae* deletion library for strains in which OspB was no longer able to inhibit yeast growth in the presence of caffeine (**Fig. 2A**). The OspB-mediated growth defect was diminished in the absence of 81 genes, including several whose gene products act downstream of TORC1 (**Table S4**). Deletion of these genes would be expected to uncouple TORC1 signaling from its downstream transcriptional response, thereby decreasing the sensitivity to TORC1 inhibitors. This validates the ability of the screen to identify host factors required for the OspB-mediated growth phenotype and confirms a role of TORC1 signaling therein.

**FIGURE 2:**
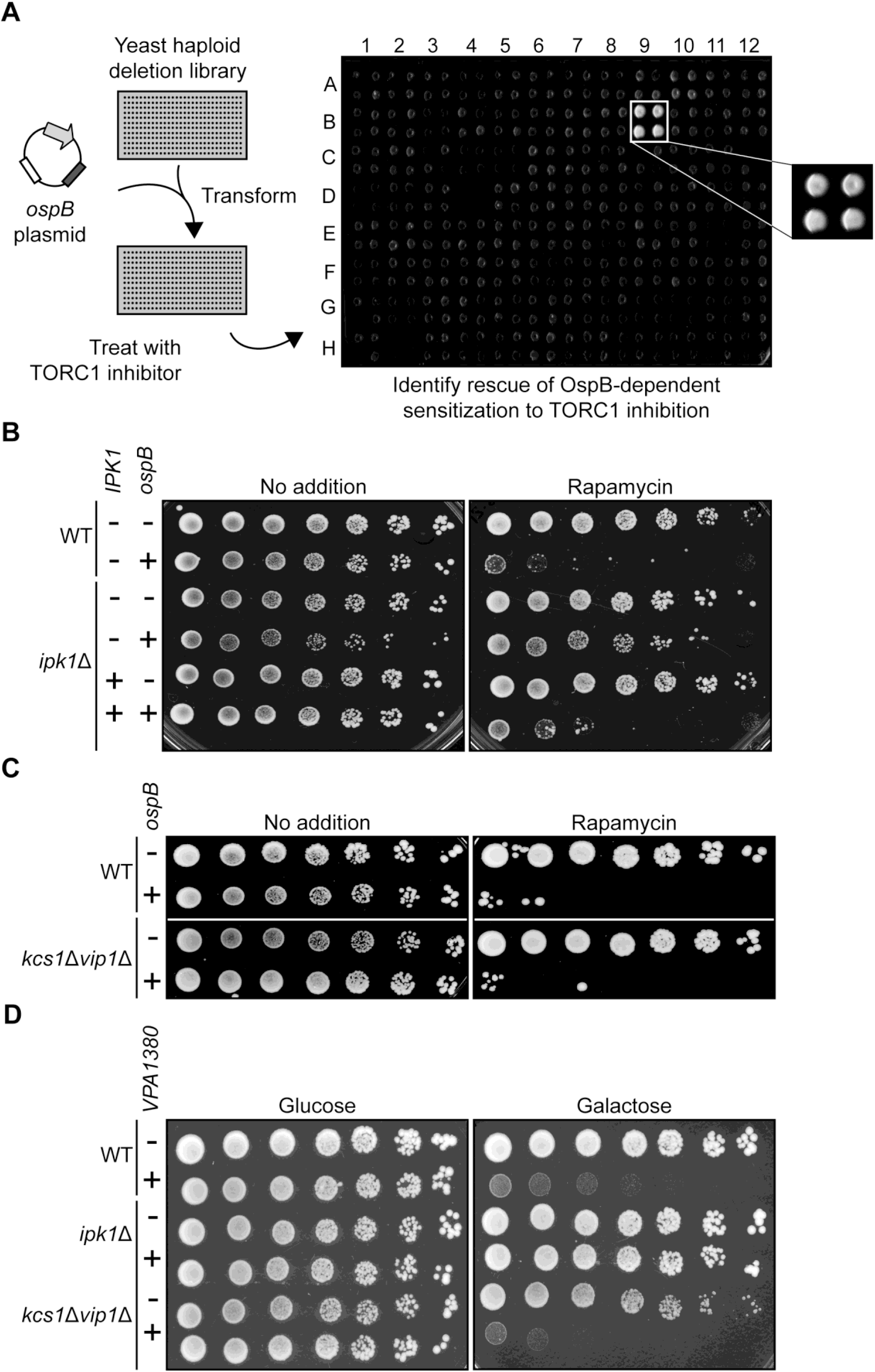
Inhibition of yeast growth by OspB-family effectors requires inositol hexakisphosphate. (A) Schematic of the deletion library screen designed to identify yeast host factors required for OspB-mediated growth inhibition in the presence of caffeine. An example of a quadruplicate-spotted output plate is shown, with one hit magnified. (B) Growth of yeast strains expressing *ospB* constructs or vector control in the presence or absence of *IPK1*. Serial dilutions were spotted on media with or without rapamycin. (C) Growth of yeast strains expressing *ospB* constructs or vector control in the presence or absence of both genes encoding the IP_6_ kinases Kcs1p and Vip1p. Serial dilutions were spotted on media with or without rapamycin. (D) Growth of yeast strains expressing *VPA1380* or vector control in the WT, *ipk1*Δ, and *kcs1*Δ*vip1*Δ backgrounds. Serial dilutions were spotted on media repressing (glucose) or inducing (galactose) *VPA1380* construct expression.

The screen also identified *IPK1*, which encodes the enzyme responsible for the generation of inositol hexakisphosphate (IP_6_), as required for yeast growth inhibition by OspB. Using an independent *ipk1* deletion strain, we confirmed that *IPK1* is required for OspB-mediated growth sensitivity to TORC1 inhibitors and found that reintroduction of *IPK1 in trans* restored growth inhibition (**Fig. 2B**). Of note, IP_6_ is an allosteric activator of the CPDs of RtxA and TcdA, and it is required for cysteine protease activity *in vitro* (Prochazkova & Satchell, 2008; Reineke et al., 2007). *In vitro* data assessing the role of IP_6_ and a more highly phosphorylated inositol pyrophosphate species (IP_7_) in the autoprocessing of TcdB indicate that IP_7_ is also a potent activator of TcdB cysteine protease activity (Savidge et al., 2011). Since an *ipk1* mutant lacks IP_6_ and all IP_7_ and IP_8_ inositol pyrophosphate species (Saiardi et al., 2002), we tested whether these inositol pyrophosphatase species were dispensable for OspB enzymatic activity by assessing growth inhibition in the absence of both yeast inositol hexakisphosphate kinases Kcs1p and Vip1p (Mulugu et al., 2007). The *kcs1*Δ*vip1*Δ mutant still displayed OspB-dependent sensitivity to TORC1 inhibitors, indicating that IP_6_ is sufficient to stimulate the activity of OspB (**Fig. 2C**). Furthermore, we confirmed previous data (Calder et al., 2014) that showed that IP_6_ is sufficient for enzymatic activation of the OspB ortholog VPA1380 from *Vibrio parahaemolyticus* (**Fig. 2D**), pointing to the Ipk1p-dependency of VPA1380 for yeast growth inhibition resulting specifically from the loss of IP_6_ rather than inositol pyrophosphate species.

### The arginine N-degron pathway is required for growth inhibition by OspB

Our deletion screen for host factors required for OspB-mediated sensitization of yeast to TORC1 inhibition also identified several components of the arginine N-degron pathway. This protein degradation pathway recognizes the neo-N-termini of polypeptides generated by cleavage or processing events and targets the polypeptides to the proteasome (Varshavsky, 2019). If the N-terminal residue of the C-terminal fragment produced upon protein cleavage is glutamine or asparagine, this destabilizes the fragment, directing its degradation *via* the arginine N-degron pathway (Gonda et al., 1989). The N-terminal Gln or Asn residue is deamidated to a Glu or Asp residue, respectively, by the N-terminal amidase Nta1p. The polypeptide is arginylated at the N-terminus by the arginine transferase Ate1p, permitting subsequent recruitment of the E3-E2 ubiquitin ligase N-recognin complex Ubr1p-Rad6p, which ubiquitinates the N-degron for degradation by the proteasome (**Fig. 3A**) (Baker & Varshavsky, 1995; Bartel et al., 1990; Dohmen et al., 1991; Richter-Ruoff et al., 1992).

**FIGURE 3:**
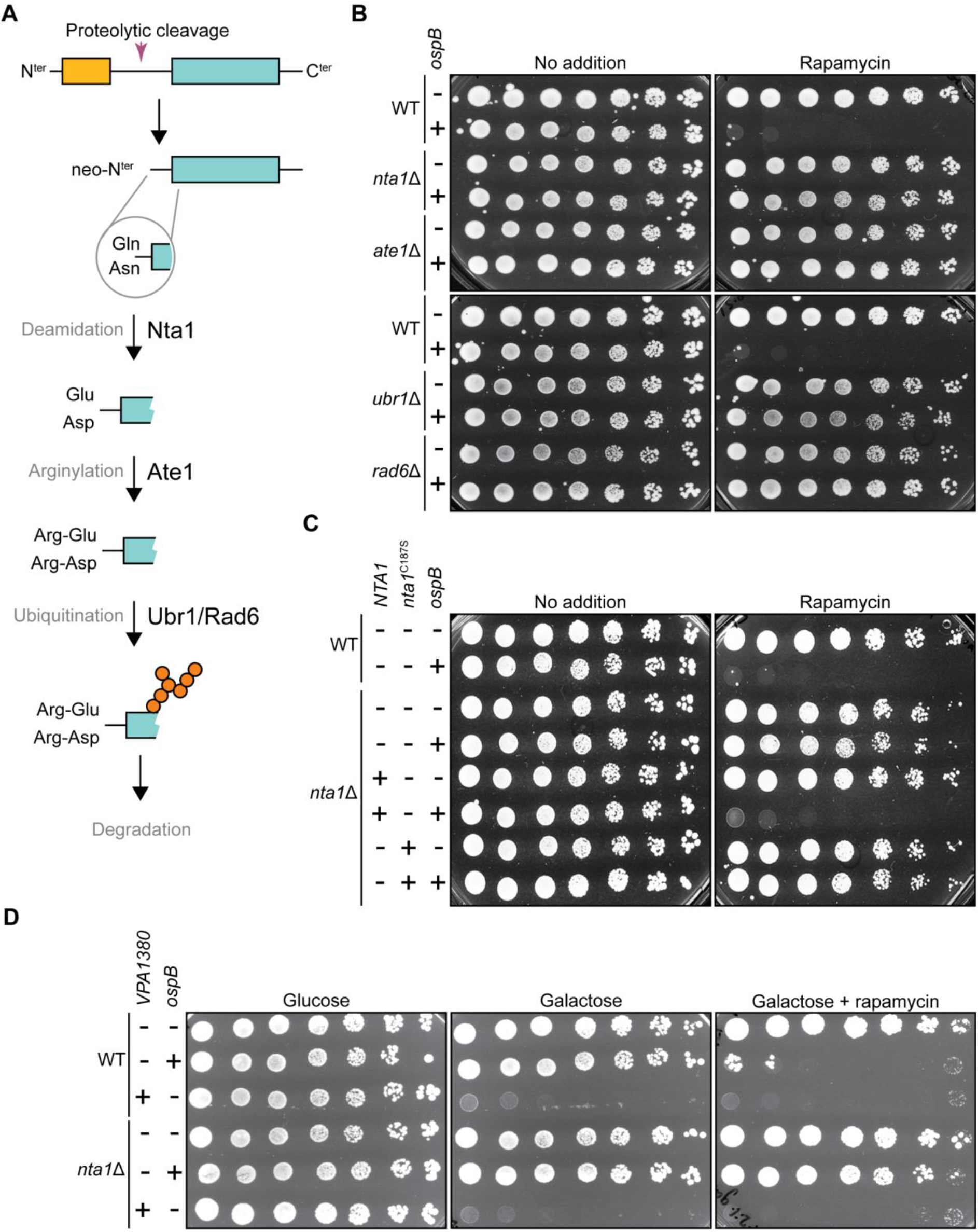
The arginine N-degron pathway is required for growth inhibition by OspB. (A) Schematic of the arginine N-degron pathway. (B) Growth of yeast strains expressing *ospB* or vector control in the presence or absence of genes encoding components of the arginine N-degron pathway. Serial dilutions were spotted on media with or without rapamycin (*n* = 3). (C) Growth of yeast strains expressing *ospB* or vector control in the presence or absence of a functional *NTA1* allele. Serial dilutions were spotted on media with or without rapamycin (*n* = 3). (D) Growth of yeast strains expressing *ospB, VPA1380*, or vector control in the presence or absence of *NTA1*. Serial dilutions were spotted on media with or without rapamycin, in conditions repressing (glucose) or inducing (galactose) *VPA1380* expression (*n* = 3).

Deletion of *nta1, ate1, ubr1*, or *rad6* each rescued the growth inhibition phenotype catalyzed by OspB (**Fig. 3B**). Moreover, introduction of a plasmid expressing *NTA1* from its native promoter complemented the *nta1* mutant, whereas complementation with a *nta1*(C187S) allele, encoding a catalytically inactive amidase, did not restore OspB-dependent sensitivity to TORC1 inhibition (**Fig. 3C** and **Fig. S3**). These results show that the host arginine N-degron pathway is required for growth inhibition by OspB and suggests that the generation of an N-degron harboring an N-terminal Gln or Asn is a necessary step in this process.

A parallel screen using a yeast over-expression library (Sopko et al., 2006) to identify suppressors of OspB-mediated sensitivity to caffeine found that expression of *BRE1* from a multicopy vector rescued the OspB-dependent growth defect (**Table S5**). Bre1p is an E3 ubiquitin ligase that monoubiquitinates histone H2B to regulate chromatin structure, in conjunction with Rad6p as the E2 ubiquitin conjugating enzyme (Hwang et al., 2003; Wood et al., 2003). We hypothesized that the mechanism of rescue by Bre1 overexpression is its sequestration of Rad6p from the arginine N-degron pathway, effectively phenocopying a *rad6* mutant (**Fig. S4A**). We found that the E2 enzymatic activity of Rad6p is required for growth inhibition by OspB, since production of a catalytically inactive Rad6p (C88S) variant did not complement a *rad6* mutant, whereas the wild type *RAD6* allele did (**Fig. S4B**). In contrast, overexpression of a catalytically dead Bre1p (C663S) variant rescued the growth defect (**Fig. S4C**), indicating that the E3 ubiquitin ligase activity of Bre1p is dispensable for suppression of the OspB phenotype, consistent with the proposed mechanism of suppression being Rad6p sequestration. Expression of an additional copy of *RAD6* negated the suppression phenotype of Bre1p overexpression (**Fig. S4D**), suggesting that higher levels of Bre1p rescue growth through indirectly reducing flux through the arginine N-degron pathway. Taken together, these data demonstrate that the activity of the arginine N-degron pathway is critical for OspB to sensitize yeast to TORC1 inhibition.

VPA1380 expression is toxic to yeast even in the absence of TORC1 inhibitors, indicative of a mechanism divergent to that of OspB. In addition, the absence of *nta1* or other arginine N-degron pathway components did not perturb the growth inhibition elicited by VPA1380, suggesting that the outcome of its activity differs from that of OspB (**Fig. 3D & Fig. S5**). Furthermore, no role was found for the formyl-methionine or proline N-degron pathways (Chen et al., 2017; J.-M. Kim et al., 2018; Melnykov et al., 2019) in VPA1380-mediated growth inhibition (**Fig. S5**). Thus, despite the homology between OspB and VPA1380, and that both T3SS effectors inhibit yeast growth, these data suggest that these bacterial proteins elicit toxicity *via* divergent mechanisms.

### OspB cleaves the TORC1 component Tco89p

Since yeast expressing OspB are sensitive to TORC1 inhibitors, we postulated that OspB either perturbs TORC1 signaling upstream of TORC1 or directly manipulates the TORC1 complex itself. Genetic ablation of individual nutrient sensing pathways upstream of TORC1 (**Fig. 4A**) (Binda et al., 2009; Dubouloz et al., 2005; Hughes Hallett et al., 2014; A. Kim & Cunningham, 2015; Yuan et al., 2017) did not rescue the OspB phenotype (**Fig. S6A**). With respect to *pib2*, which encodes a glutamine sensor that activates TORC1 in parallel to the amino acid-responsive Gtr1p/Gtr2p pathway (Tanigawa et al., 2021; Ukai et al., 2018), a yeast strain constitutively producing OspB in the absence of *pib2* could not be generated; however, expression of *ospB* from a galactose inducible promoter completely inhibited the growth of a *pib2* mutant without the need for low levels of caffeine or rapamycin (**Fig. 4B**). Genetic ablation of components of TORC1 or the Gtr1p/Gtr2p branch of amino acid sensing is synthetically lethal in a *pib2*Δ background (A. Kim & Cunningham, 2015), and we find that OspB is still toxic in a *gtr1*Δ*gtr2*Δ double mutant (**Fig. S6B**). We therefore conclude that OspB likely targets the TORC1 complex itself.

**FIGURE 4:**
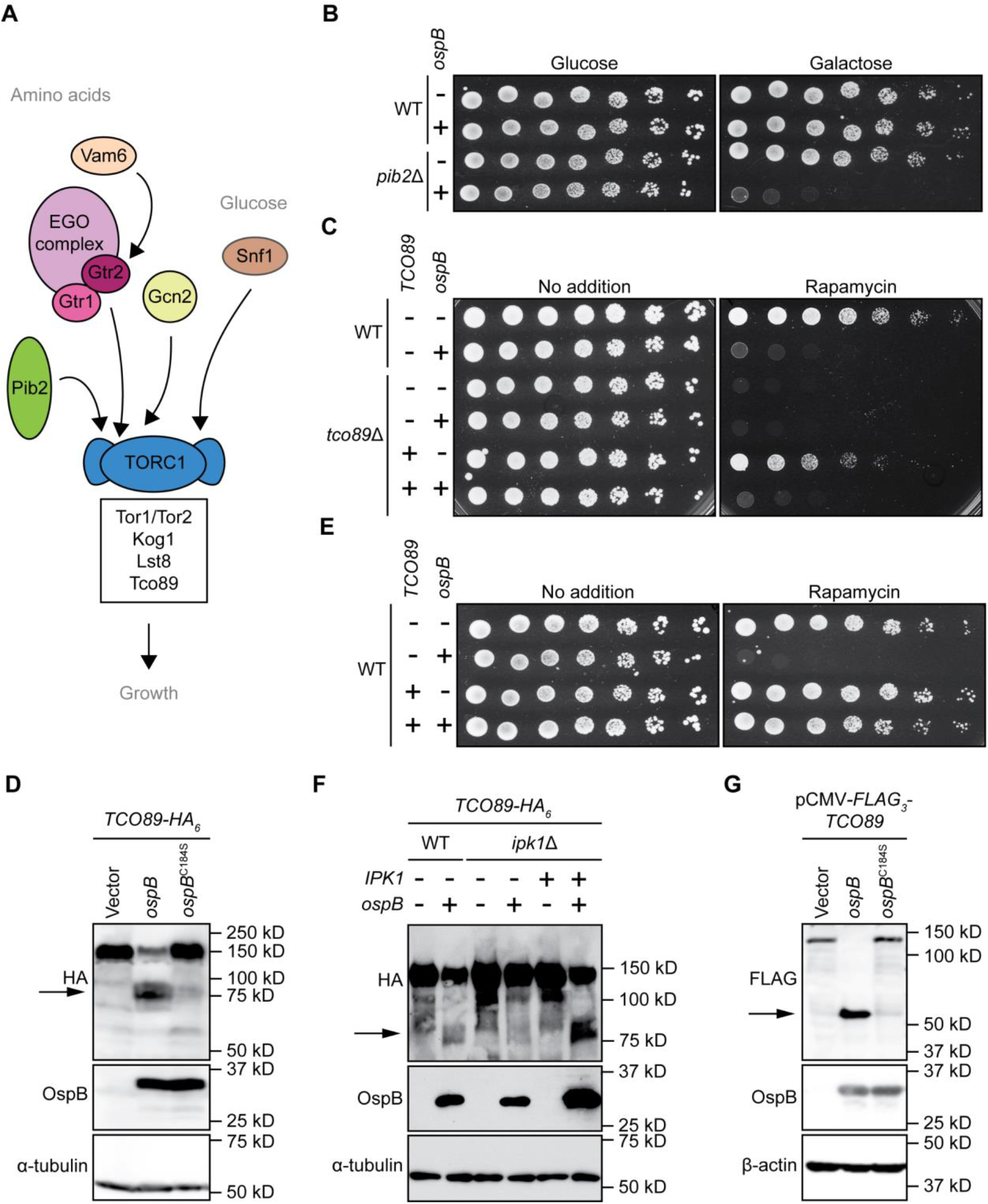
The TORC1 subunit Tco89p is cleaved by OspB. (A) Diagram of the yeast TORC1 signaling network. (B) Growth of yeast strains expressing *ospB* or vector control in the presence or absence of *PIB2*. Serial dilutions were spotted on media containing glucose (repressing *ospB* expression) or galactose (inducing *ospB* expression) (*n* = 3). (C) Growth of yeast strains expressing *ospB* or vector control in the presence or absence of *tco89*. Serial dilutions were spotted on media with or without rapamycin (*n* = 4). (D) Western blot assessing cleavage of Tco89p in yeast, in the presence of OspB, OspB(C184S) or vector control. Alpha tubulin is the loading control. The arrow marks the Tco89p C-terminal cleavage product (*n* = 4). (E) Growth of wild type yeast expressing *ospB* or vector control, in the presence or absence of *TCO89* expression from a multi-copy plasmid. Serial dilutions were spotted on media with or without rapamycin (*n* = 3). (F) Western blot assessing cleavage of Tco89p by OspB in wild type or *ipk1* mutant yeast. Alpha tubulin is the loading control. The arrow marks the Tco89p C-terminal cleavage product (*n* = 3). (G) Western blot assessing cleavage of a Tco89p construct expressed in HEK293T cells, in the presence of OspB, OspB(C184S) or vector control. Beta-actin is the loading control. The arrow marks the Tco89p N-terminal cleavage product (*n* = 3).

We tested whether any of the four proteins that comprise the TORC1 complex - Kog1p, Lst8p, Tco89p and Tor1p/Tor2p (Reinke et al., 2004) – are perturbed by OspB activity. Among these four proteins, only Tor1p and Tco89p are non-essential. We found that neither the essential TORC1 components nor Tor1p is cleaved by OspB (**Fig. S7A**). Since deletion of *tco89* rendered yeast hypersensitive to rapamycin (Reinke et al., 2004), it was not possible to test for an effect of OspB in this growth assay. OspB-dependent sensitivity to TORC1 inhibitors was restored upon *TCO89* complementation (**Fig. 4C**).

Assessment of Tco89p abundance revealed that in the presence of OspB, full length Tco89p levels were decreased and a faster migrating Tco89p band that was recognized by an antibody to the C-terminal tag was present (**Fig. 4D**). This faster migrating band was not observed in the presence of the catalytically inactive OspB C184S mutant, indicating that its generation depends on OspB catalytic activity and that the faster migrating band is a C-terminal Tco89p cleavage product. Overexpression of *TCO89* in wild type yeast rescued the OspB-dependent growth defect (**Fig. 4E**), presumably because resulting increase in Tco89p levels likely raises the threshold at which sensitivity to TORC1 inhibitors occurs.

Cleavage of endogenous Tco89p was abolished in an *ipk1* background and restored by complementation of *IPK1*, indicating that requirements for the yeast growth phenotype are associated with the Tco89p cleavage phenotype (**Fig. 4F**). Tco89p cleavage by OspB was unaffected in a *kcs1*Δ*vip1*Δ mutant (**Fig. S7B**), providing additional evidence that IP_6_ is the inositol phosphate species acting as the cofactor for OspB protease activity. Transfection of mammalian cells with both OspB and Tco89p revealed that OspB cleavage of Tco89p occurs (**Fig. 4G**), which indicates that OspB functions as a protease in mammalian cells and suggests that Tco89p may be a direct substrate of OspB protease activity.

Processing of Tco89p by VPA1380 was not observed (**Fig. S8**), further supporting that the mechanisms of OspB and VPA1380 are divergent. Together, these data demonstrate that, upon its allosteric activation by IP_6_, OspB cleaves the TORC1 component Tco89p triggering sensitivity to inhibition of TORC1 signaling.

We assessed the role of the arginine N-degron pathway in the stability of the C-terminal Tco89p cleavage product generated by OspB. Treatment of yeast with the proteasome inhibitor MG-132 increased the abundance of the Tco89p C-terminal fragment, indicating that this fragment is a substrate of the proteasome (**Fig. 5A**). Deletion of *nta1* resulted in an increase in the abundance of the Tco89p C-terminal fragment (**Fig. 5B**), and its levels were restored by reintroduction of *NTA1*, but not by reintroduction of the C187S inactive amidase mutant, indicating that the C-terminal fragment of Tco89p is the N-degron that mediates growth inhibition caused by the protease activity of OspB.

**FIGURE 5:**
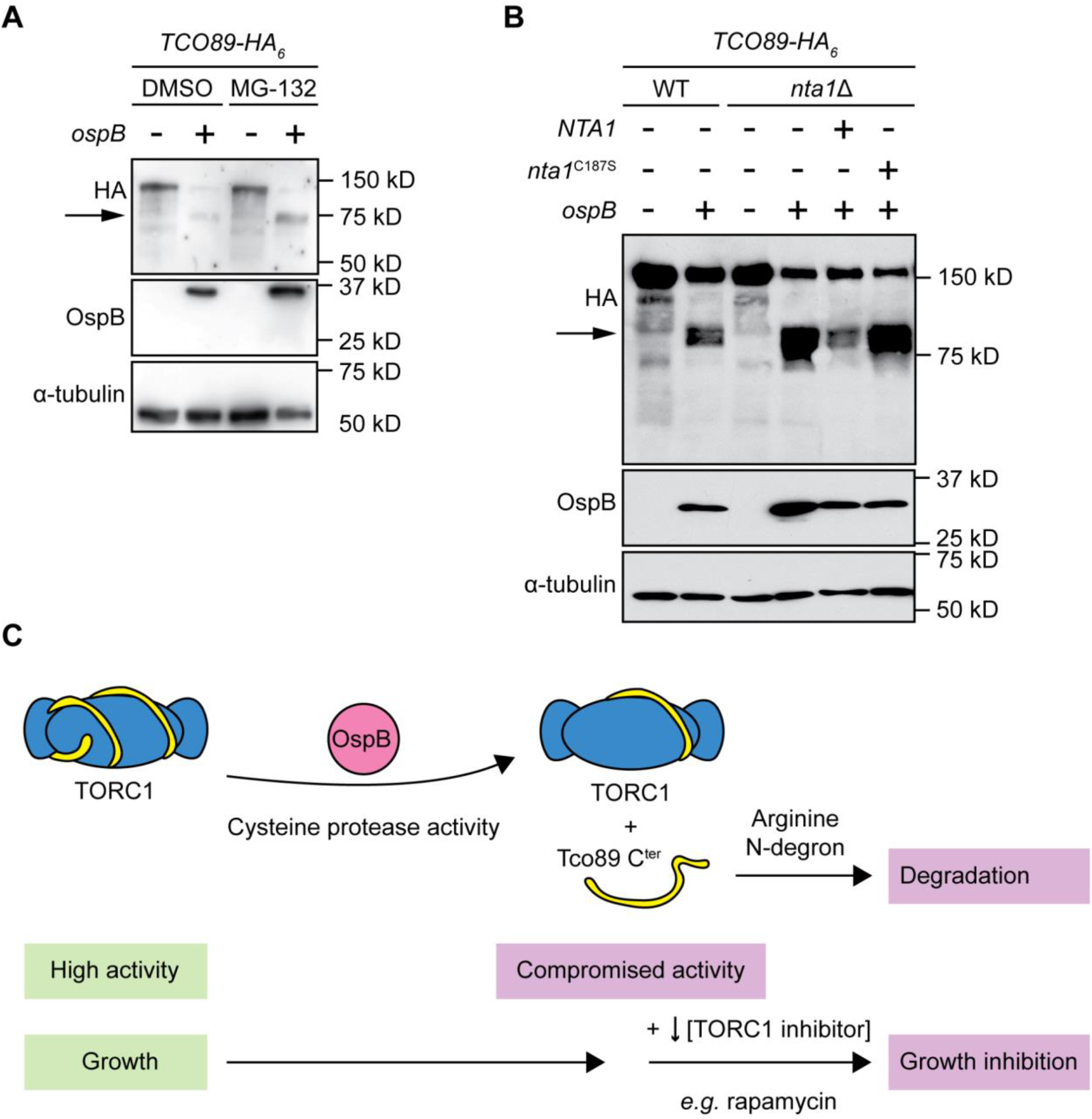
Degradation of the Tco89p C-terminal fragment by the arginine N-degron pathway. (A) Western blot assessing cleavage of Tco89p by OspB in the presence and absence of proteasome inhibitor MG-132. Alpha tubulin acts as a loading control. The arrow marks the Tco89p C-terminal cleavage product (*n* = 3). (B) Western blot assessing cleavage of Tco89p by OspB in the presence or absence of a functional *NTA1* allele. Alpha tubulin is the loading control. The arrow marks the Tco89p C-terminal cleavage product (*n* = 3). (C) Model of the mechanism of OspB-mediated sensitization of yeast to TORC1 inhibition.

In summary, we found that the *Shigella* T3SS effector OspB is a cysteine protease that cleaves the TORC1 component Tco89p, thereby generating an N-degron, and that the N-degron is targeted for degradation by the arginine N-degron pathway (**Fig. 5C**). Cleavage of Tco89p by OspB and host-mediated degradation of the C-terminal fragment is responsible for sensitization of the TORC1 complex to inhibition and the associated inhibition of yeast growth.

## Discussion

The evidence presented here collectively demonstrates that the *Shigella* T3SS effector OspB is a cysteine protease and that it requires inositol hexakisphosphate for its activity. OspB is required for cleavage of Tco89p, a component of yeast TORC1, and upon expression of the two proteins in mammalian cells, OspB is sufficient to cleave Tco89p (**Fig. 4**), indicating that OspB mediated cleavage of Tco89p is either direct or depends on factors that are conserved between yeast and mammalian cells. OspB is structurally homologous to the cysteine protease domains of the bacterial MARTX toxins TcdA, TcdB and RtxA, with conservation of the catalytic cysteine and histidine residues (**Fig. 1 and S2**) (Egerer et al., 2007; Lupardus et al., 2008; Reineke et al., 2007; Shen et al., 2009). In OspB, the conserved cysteine and histidine are each required for both the OspB growth inhibition phenotype in yeast and Tco89p cleavage in yeast and mammalian cells (**Fig. 4** and data not shown).

Like TcdA, TcdB and RtxA, OspB requires inositol hexakisphosphate for its activity (**Fig. 2** and **4**) (Egerer & Satchell, 2010). Using a genetic approach, we exclude inositol pyrophosphate species as being necessary for OspB activity and define the inositol phosphate species requirement as IP_6_; however, it is possible that an inositol pyrophosphate species may be sufficient for stimulating activity *in vitro*, as found for IP_7_ and *C. difficile* TcdB (Savidge et al., 2011). The requirement for a host-specific cofactor, such as IP_6_, ubiquitin, calmodulin, or cyclophilins, for the activation bacterial effectors is increasingly appreciated (Anderson et al., 2011; Coaker et al., 2005; Drum et al., 2002; Mittal et al., 2010; Sreelatha et al., 2020; Tyson & Hauser, 2013) and necessarily restricts enzymatic activity to the context of host infection. Together, these findings provide strong evidence that OspB is a cysteine protease in the family of proteases represented by the cysteine protease domains of the MARTX toxins and large clostridial cytotoxins.

Two other *Shigella* T3SS effectors, IpaJ and OspD3, are also cysteine proteases. IpaJ and OspD3 are divergent from OspB and their substrates are distinct, constituting Rho GTPases and necroptotic signaling factors, respectively (Ashida et al., 2020; Burnaevskiy et al., 2015). Genes encoding T3SS effector proteins are often acquired by horizontal gene transfer (Brown & Finlay, 2011), thus homologous effectors are commonly secreted by the T3SSs of pathogens displaying similar host tropism. The T3SS2 effector of *V. parahaemolyticus* VPA1380 is homologous to OspB, yet we find that it likely targets a different host substrate. First, yeast growth inhibition by OspB requires reduction of TORC1 activity by either chemical or genetic intervention (**Fig. 1** and **4**), whereas VPA1380 is toxic to yeast in the absence of additional stressors (**Fig. 2**) (Calder et al., 2014). Second, we find that VPA1380 neither cleaves Tco89p nor generates a substrate cleavage product that requires the arginine N-degron pathway for its degradation (**Fig. 3**, **S5**, and **S8**).

OspB cleaves a component of TORC1 to produce a C-terminal fragment that enters the arginine N-degron pathway for proteasomal degradation, rendering the cells hypersensitive to TOR inhibition. Tco89p cleavage and degradation appears to be entirely responsible for the TOR inhibitor hypersensitivity phenotype stimulated by OspB, as degradation of the C-terminal fragment phenocopies a *tco89*Δ mutant growth defect in the presence of either rapamycin or caffeine (**Fig. 4C**) (Reinke et al., 2004, 2006; Slagowski et al., 2008). Consequently, complementation with multi-copy Tco89p is associated with reduced sensitivity to inhibition of TORC1 by a combination of OspB and chemical inhibitors (**Fig. 4E**). The dependence on *NTA1*, the upstream-most enzyme in the arginine N-degron pathway (**Fig. 3**), indicates that OspB cleavage results in a tertiary arginine N-degron, with glutamine or asparagine at the neo-N-terminus of the C-terminal Tco89p cleavage product, since deamidation of the product by Nta1p is a critical step in its degradation (Gonda et al., 1989).

The migration of Tco89p constructs in SDS-PAGE is slower (at around 150 kD) than expected for the 89 kD protein (**Fig. 4**). We hypothesize that the retarded migration of Tco89p is due to significant phosphorylation by the TORC1 kinase (Huber et al., 2009; Oliveira et al., 2015). Irrespective of the cause, prediction of the OspB cleavage site producing the C-terminal fragment cannot be based on gel migration. Tco89p is an intrinsically disordered protein and these proteins are often enriched in phosphorylation sites (Miao et al., 2018). Moreover, post-translational modification is a frequent regulator of intrinsically disordered proteins, so it is conceivable that TORC1 regulates its own function by altering the phosphorylation state of Tco89p, consistent with a role for this protein in formation of inhibitory TORC1 “body” formation during glucose and nitrogen starvation (Hughes Hallett et al., 2015; Sullivan et al., 2019).

Of note, there is no obvious homolog of Tco89p in mammalian cells. However, since Tco89p is intrinsically disordered, due to the absence of structural constraints, it would be expected to have evolved rapidly and to have undergone positive selection at specific sites, resulting in the acquisition of new functions (Afanasyeva et al., 2018), leading us to postulate that the mammalian functional homolog is divergent at the sequence level. Notwithstanding this potential lack of recognizable sequence identity, our yeast OspB phenotype of sensitization to TOR inhibition is similar to our prior finding of OspB mediated sensitization to rapamycin in fibroblasts (Lu et al., 2015), bolstering the relevance of the yeast model.

The potential utility of identifying a substrate of a microbial protease in a heterologous system, as we did here for OspB, is exemplified by the work leading to the identification of the physiological ligand of the NLRP1 inflammasome. The *Bacillus anthracis* lethal factor protease efficiently cleaves a disordered linker in murine NLRP1B and rat NLRP1, releasing an arginine N-degron, degradation of which led to inflammasome activation in macrophages and pyroptotic cell death (Boyden & Dietrich, 2006; Levinsohn et al., 2012; Wickliffe et al., 2008). Anthrax is primarily a pathogen of humans, and lethal factor does not cleave the human NLRP1 homolog (Chavarría-Smith et al., 2016). Yet, these studies facilitated the recent determination that dependence on functional degradation is a conserved feature of NLRP1 activation (Chui et al., 2019; Sandstrom et al., 2019; Xu et al., 2019), and the subsequent molecular identification of enteroviral proteases as the physiological activators of the human NLRP1 inflammasome, in which cleavage of the aforementioned disordered linker generates a glycine N-degron (Robinson et al., 2020; Tsu et al., 2021). By analogy, through determination of the activity of OspB, our study provides an important insight into its substrate specificity and phenotypic impact, which will facilitate identification of mammalian substrates.

## Acknowledgments

We thank Austin C. Hachey and Yang Fu for technical assistance, and Ted Powers and Nick Laribee for the gift of yeast strains. The work was supported by Department of Defense grant TS160046 (to M.B.G.), funding from the Massachusetts General Hospital Executive Committee on Research (to M.B.G.), NIH grants T32 AI007061 and F32 AI131582 (to H.D.E.), and R01 AI064285 (to C.F.L.). The authors have no conflicts of interest.

## Materials and Methods

### Strains and media

All strains, plasmids and primers are listed in **Table S1**, **S2** and **S3**, respectively. *E. coli* DH10B (Grant et al., 1990) was used as the routine cloning host and was grown in Luria broth at 37 °C with agitation. *S. cerevisiae* S288C was used as the heterologous expression host to probe the roles of host proteins in the function of OspB and was routinely cultured at 30 °C in yeast extract-peptone-dextrose (YPD) broth or in synthetic selective media (MP Biomedicals) lacking histidine, uracil and/or leucine for auxotrophic selection. 1.5 % (w/v) agar was added for solid media formulations, and where appropriate, media was supplemented with 50 μg/ml ampicillin (Sigma, A9518), 2% (w/v) D-glucose (Fisher Scientific, D16-10), 2% D-(+)-raffinose (Sigma, R7630), 2% (w/v) D-galactose (VWR, 200001-176), 300 μg/ml hygromycin (Gibco, 10687010), 200 μg/ml geneticin (Sigma, A1720). For TORC1 inhibition, solid media was supplemented with 6 mM caffeine (Sigma, C0750) or 5 nM rapamycin (Sigma, 553211) unless stated otherwise. For proteasome inhibition, media was supplemented with 75 μM MG-132 (Selleck Chemicals, S2619) and 0.003% SDS. Yeast strains were transformed using the standard lithium acetate method.

### Bioinformatic analyses

*In silico* modelling of the tertiary structure of OspB was conducted on the Phyre2 server (Kelley et al., 2015), whereas alignment with the crystal structures of RtxA^VC^ (Lupardus et al., 2008) and TcdA (Pruitt et al., 2009) was achieved using the CEAlign algorithm within PyMol (Schrödinger, LLC). Protein sequences were retrieved from the non-redundant NCBI database and aligned using MUSCLE (Edgar, 2004) before manual curation to select the regions of interest.

### Yeast growth assays and protein extraction

Individual yeast transformants that constitutively express *ospB* or derivatives containing point mutations were grown in synthetic selective liquid media containing 2% D-glucose. To investigate the impact of OspB constructs on growth, yeast cells were washed and serially diluted four-fold in phosphate-buffered saline before 5 μl of each dilution were spotted on synthetic selective solid media with additives as appropriate. Assessment of protein production was done from liquid cultures. Here, subcultures were inoculated at OD_600_ 2.0 from overnight cultures and grown for two hours before harvesting for SDS-PAGE analysis using the alkaline lysis method (Kushnirov, 2000). Where construct induction was required, yeast strains were subcultured in 2% raffinose for 2 h, before supplementation with 2% galactose. Samples were harvested after four hours of protein expression. For proteasome inhibition, yeast strains were subscultured in glucose for 2 h before treatment with MG-132 or DMSO control for 3 h.

### Yeast library screening

To screen for suppressors of OspB-mediated toxicity in *S. cerevisiae* by yeast gene over-expression, the strain BY4742 pAG413GPD-*ospB* was mated with the haploid GST-fusion yeast over-expression library (Dharmacon, YSC4423) on YPD. The resulting diploids were selected by plating on non-inducing synthetic selective media containing 2% D-glucose. The screen was conducted by spotting in quadruplicate on inducing synthetic selective solid media containing 2% D-galactose and 6 mM caffeine. All steps in the screen were conducted in an automated manner as described previously (Slagowski et al., 2008). Suppressors were classified as strains which displayed qualitatively moderate to robust growth of all four spots on the caffeine plate four days after pinning. To screen for *S. cerevisiae* host factors required for OspB-dependent growth inhibition, we screened the MAT**a** haploid deletion library (Horizon, YSC1053) as previously described (Kramer et al., 2007), but with transformation of the plasmid pAG413GAL-*ospB* and assessment of growth on synthetic selective solid media containing 2% D-galactose and 6 mM caffeine after 3 days.

### Cell culture and transfection

HEK293T (ATCC) and mouse embryonic fibroblast cells (Lu et al., 2015) were maintained in Dulbecco’s modified Eagle medium (DMEM) (Gibco) supplemented with 10% (v/v) fetal bovine serum at 37 °C with 5% CO_2_. Cells were transfected with plasmids using FuGENE6 (Promega) according to the manufacturer’s instructions, and experimental samples were analyzed 24-48 h after transfection.

### SDS-PAGE and immunoblotting

For immunoblot analysis, protein samples were separated by SDS-PAGE, transferred to nitrocellulose membranes and detected by western blot analysis using standard procedures. The antibodies used were peroxidase-conjugated anti-β-actin (Sigma, A3854; diluted to 1:10 000), anti-α-tubulin (Santa Cruz, sc-53030; diluted to 1:1000), anti-FLAG (Sigma, F3165; diluted to 1:1000), anti-myc (EMD Millipore 05-724; diluted to 1:1000), anti-HA (Biolegend, 901501, diluted to 1:1000) and anti-OspB (diluted to 1:10 000). The rabbit anti-OspB antibody was generated (Covance Inc.) against a 14-mer peptide of OspB located 18 residues from the C-terminus.

