## Supplementary Material for "The *Shigella* type III effector protein OspB is a cysteine protease"

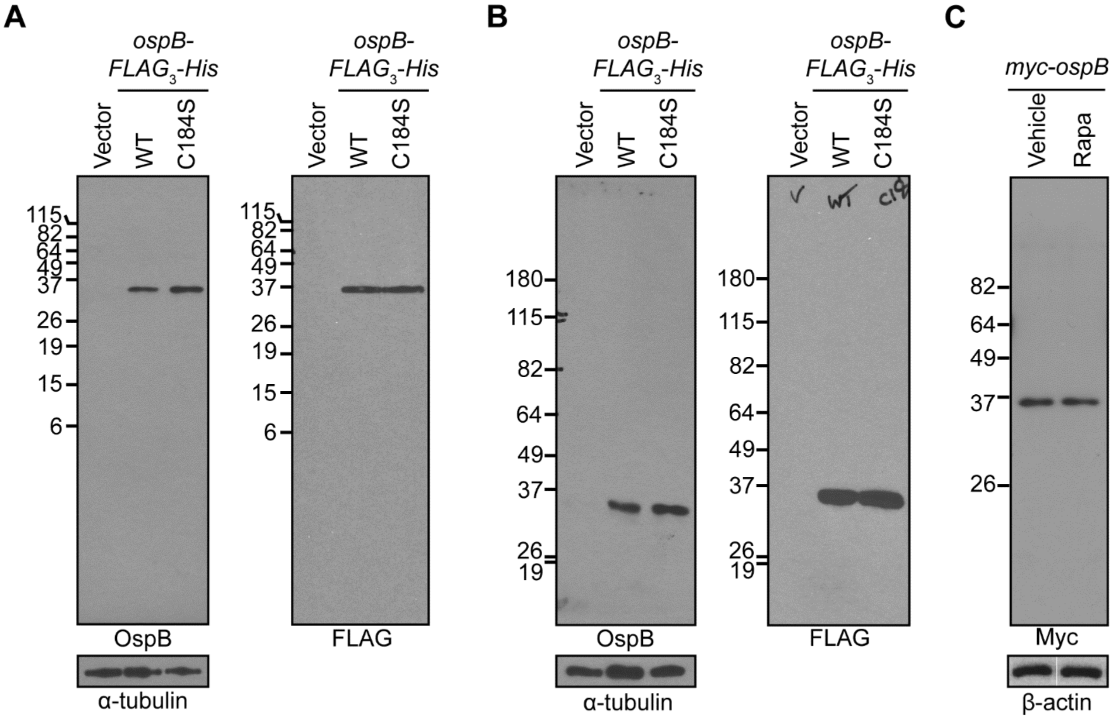


**FIGURE S1: Absence of evidence of processing of OspB in cell lysates.**

(A) Wild type OspB and OspB (C184S) expressed in yeast, detected by immunoblotting with anti-OspB and anti-FLAG antibodies after separation on a 15% SDS-PAGE gel. α-tubulin serves as a loading control.

(B) Samples from panel (A) separated on a 7.5% SDS-PAGE gel and probed as in panel (A).

(C) Transfection of mouse embryonic fibroblasts with pCMV-*myc*-*ospB*. Cells were treated with rapamycin (10 nM) (rapa) or a DMSO vehicle control. Myc-OspB detected by immunoblotting with an anti-myc antibody after separation on a 10% SDS-PAGE gel. β-actin serves as the loading control; bands from a single blot.

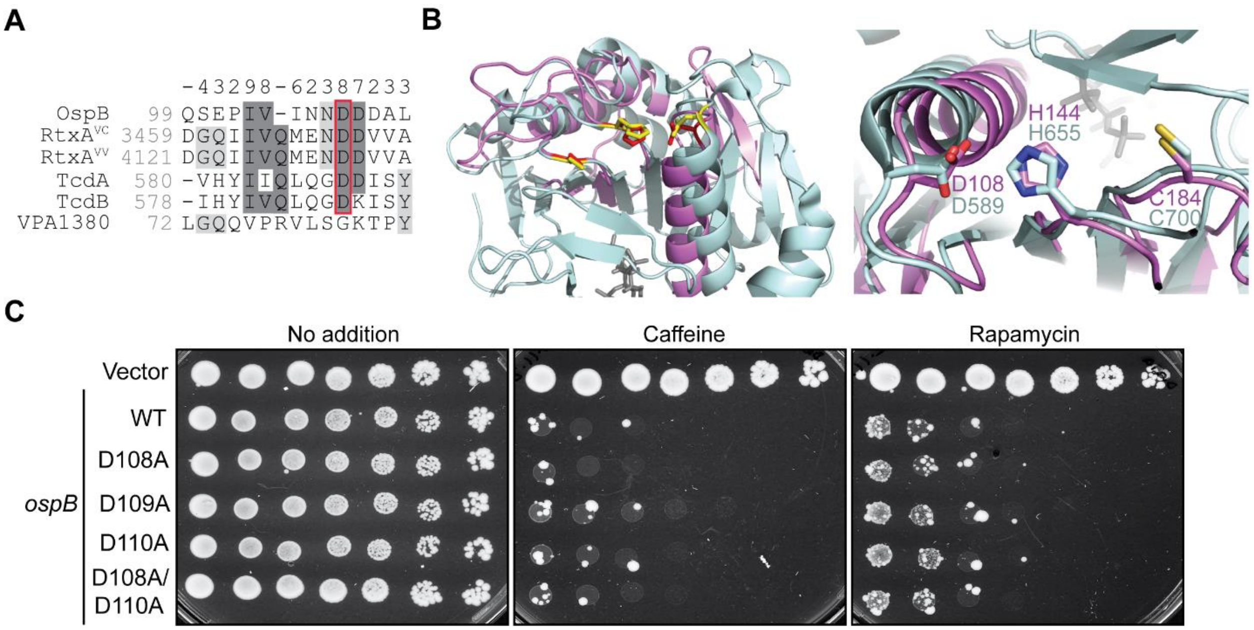


**FIGURE S2: The aspartic acid residues Asp108, Asp109, and Asp110 are not required for OspB activity.**

(A) Multiple sequence alignment of OspB with the aspartic acid residue of the cysteine protease domains of RtxA from *V. cholerae* (RtxA$^{VC}$) and *V. vulnificus* (RtxA$^{VV}$), *C. difficile* TcdA and TcdB, that in some cases stabilizes the catalytic histidine residue. The OspB ortholog VPA1380 from *V. parahaemolyticus* is also shown. The red box indicates the aspartic acid residue in the CPDs. Darkness of gray shading reflects the conservation of individual residues, and the numbers above the alignment score the conservation at each position. Asterisks denote full conservation among the aligned sequences.

(B) Cartoon depiction of a tertiary structure model of OspB (violet) on the CPD of TcdA (PDB: 3HO6) (pale cyan). In the left panel, the catalytic residues of the cysteine protease domain are denoted by yellow sticks, with the putative catalytic residues of OspB shown as red sticks. The inositol hexakisphosphate cofactor in the TcdA cysteine protease domain structure is shown in dark gray. In the right panel, an enlarged and rotated view shows the active site, highlighting the superposition of the putative OspB catalytic residues with those of the cysteine protease domain, labelled according to the color of the cartoon.

(C) Growth of yeast strains expressing *ospB* constructs or an empty vector control. Serial dilutions were spotted on media without additives or supplemented with the TORC1 inhibitors caffeine or rapamycin ($n$ = 3).

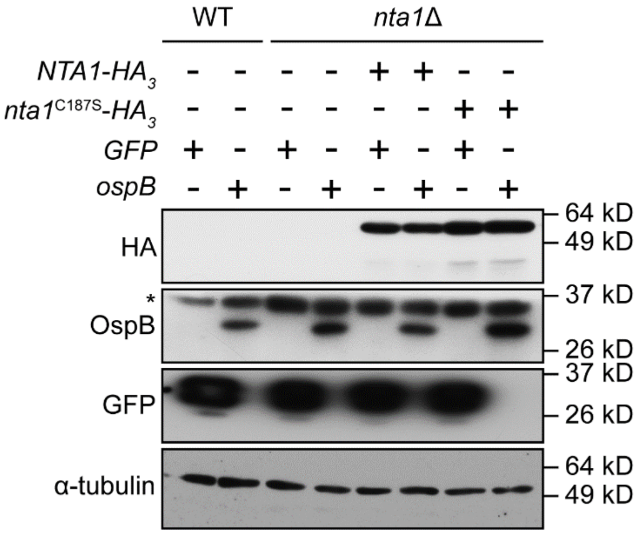


**FIGURE S3: Production of Nta1p variants.**

Western blot showing the production of the WT and catalytically inactive Nta1p variant. The asterisk marks a non-specific protein recognized by the anti-OspB antibody. α-tubulin is a loading control ($n$ = 3).

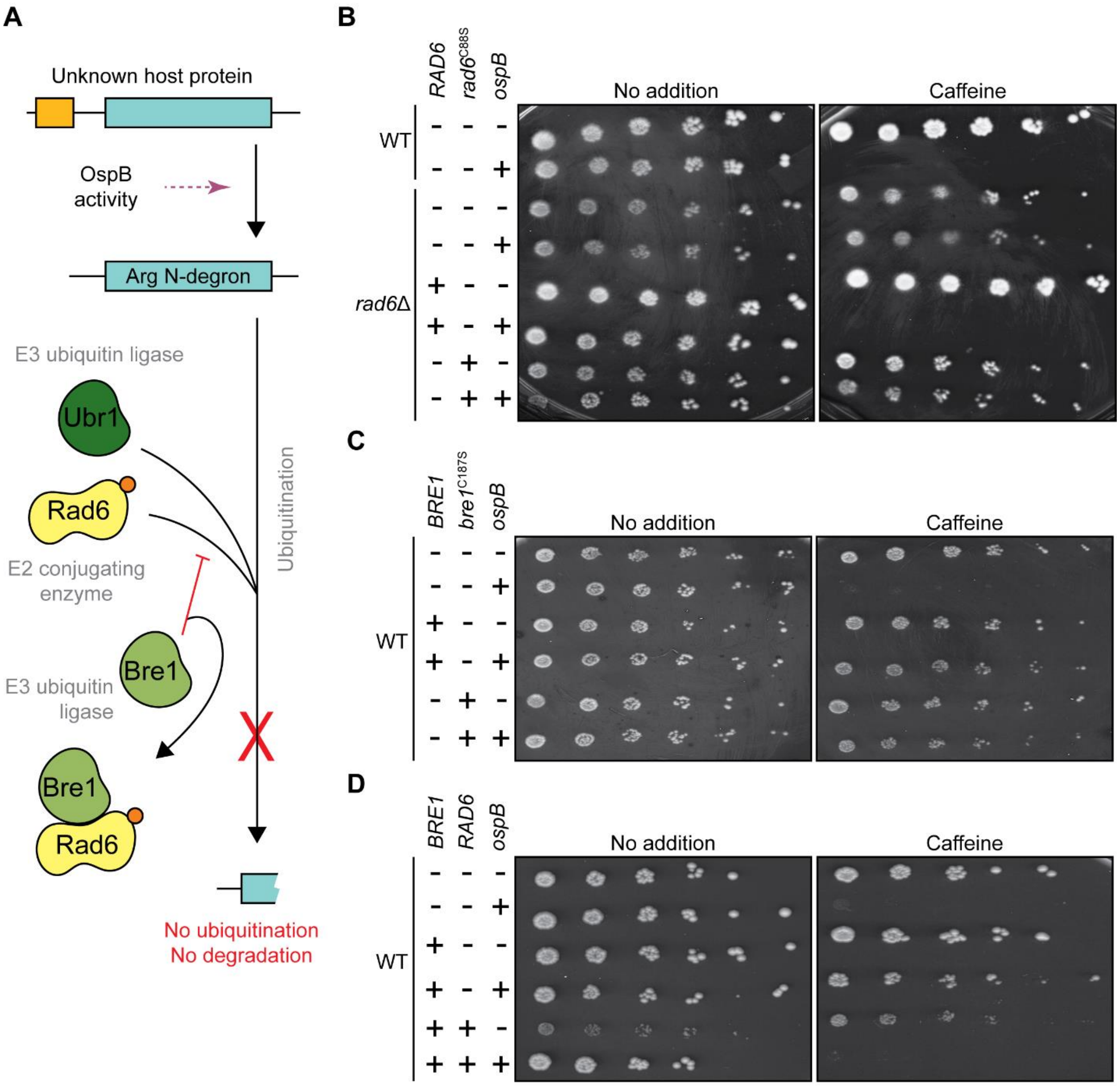


**FIGURE S4: Bre1p rescues growth inhibition through sequestration of arginine N-degron pathway component Rad6p.**

(A) Schematic of the proposed mechanism of Bre1-mediated suppression of OspB-dependent growth inhibition.

(B) Growth of yeast strains expressing *ospB* or vector control in the presence or absence of a functional *RAD6* allele. Serial dilutions were spotted on media with or without caffeine ($n$ = 3).

(C) Growth of yeast strains expressing *ospB* or vector control in the presence or absence of multi-copy *BRE1* alleles. Serial dilutions were spotted on media with or without caffeine ($n$ = 3).

(D) Growth of wild type yeast expressing *ospB* or vector control in the presence or absence of additional *BRE1* and *RAD6* gene copies. Serial dilutions were spotted on media with or without caffeine (*n* = 3).

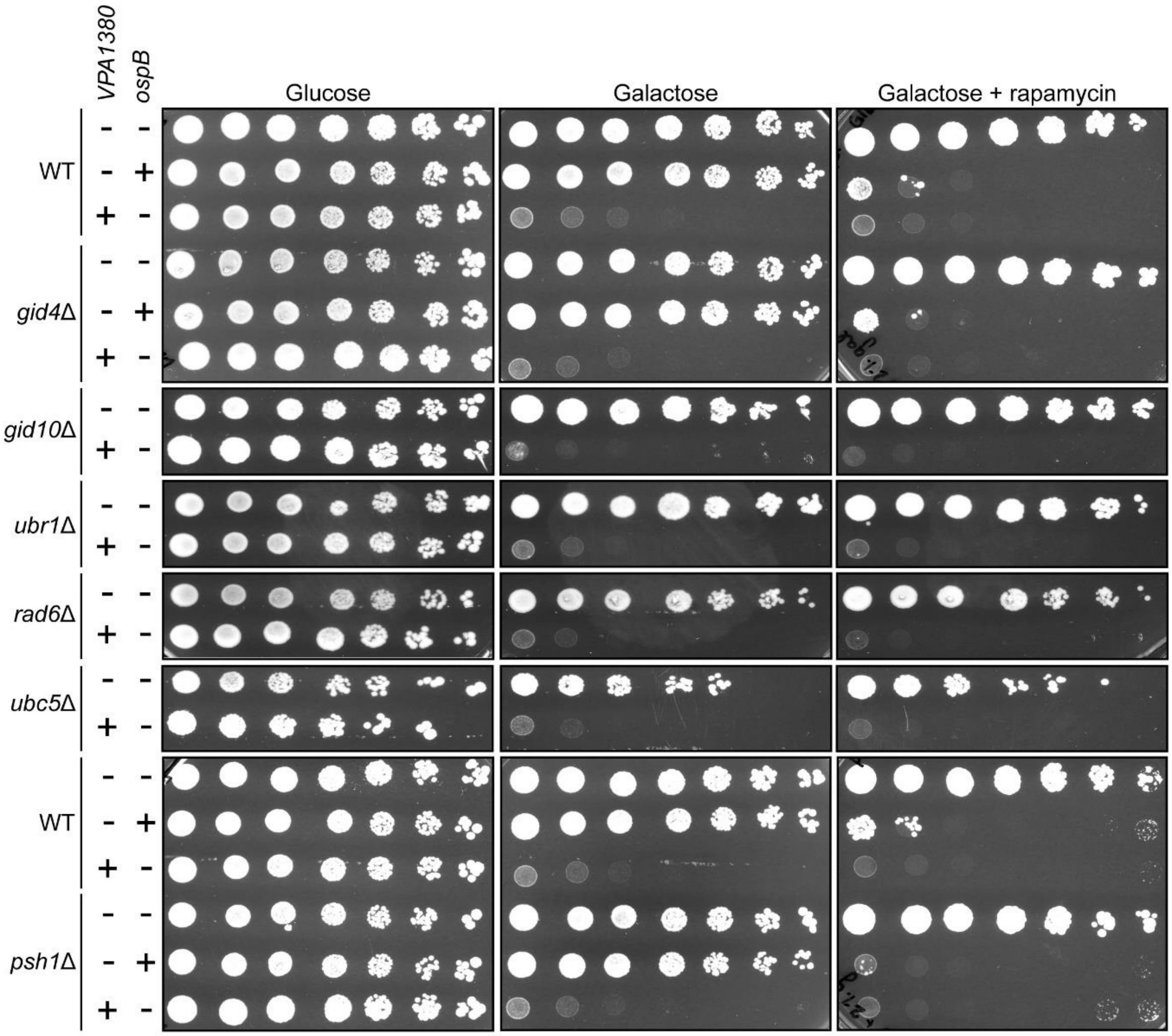


**FIGURE S5: N-degron pathways are not required for VPA1380 toxicity phenotype.**

The effect of *ospB* and *VPA1380* expression in wild type yeast, proline N-degron pathway mutants (lacking N-recognins Gid4 or Gid10), arginine N-degron pathway mutants (lacking amidase Nta1, E3 ubiquitin ligase Ubr1, or E2 conjugating enzymes Ubc5 or Rad6), or a formyl-methionine N-degron pathway mutant (lacking E3 ubiquitin ligase Psh1). Serial dilutions were spotted on media repressing (glucose) or inducing (galactose) construct expression. Rapamycin is also included as a supplement in galactose plates (*n* = 2).

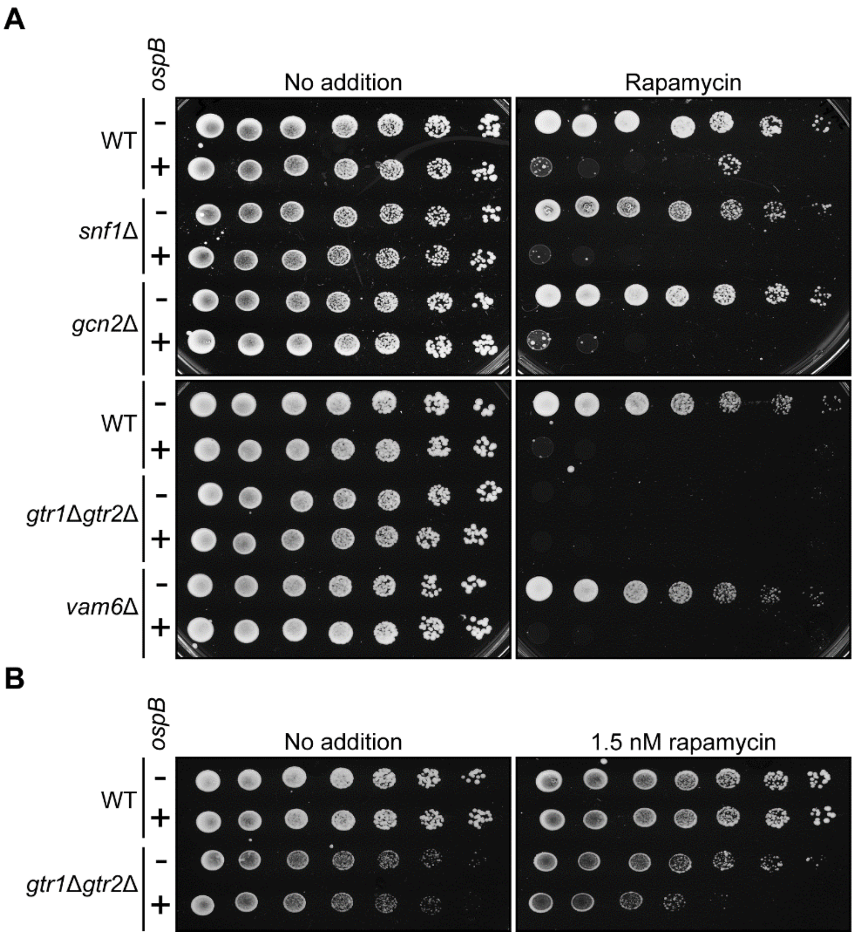


**FIGURE S6: OspB does not act upstream of TORC1.**

(A) Effect of *ospB* expression on the growth of yeast strains lacking genes involved in nutrient sensing upstream of TORC1 signaling. Serial dilutions were spotted on media with or without rapamycin (*n* = 3).

(B) Growth of wild type and a *gtr*1Δ*gtr*2Δ yeast strain expressing *ospB*, plated on solid media containing no additive or 1.5 nM rapamycin. This reduced rapamycin concentration was used due to sensitivity of the *gtr*1Δ*gtr*2Δ mutant to this compound (*n* = 3).

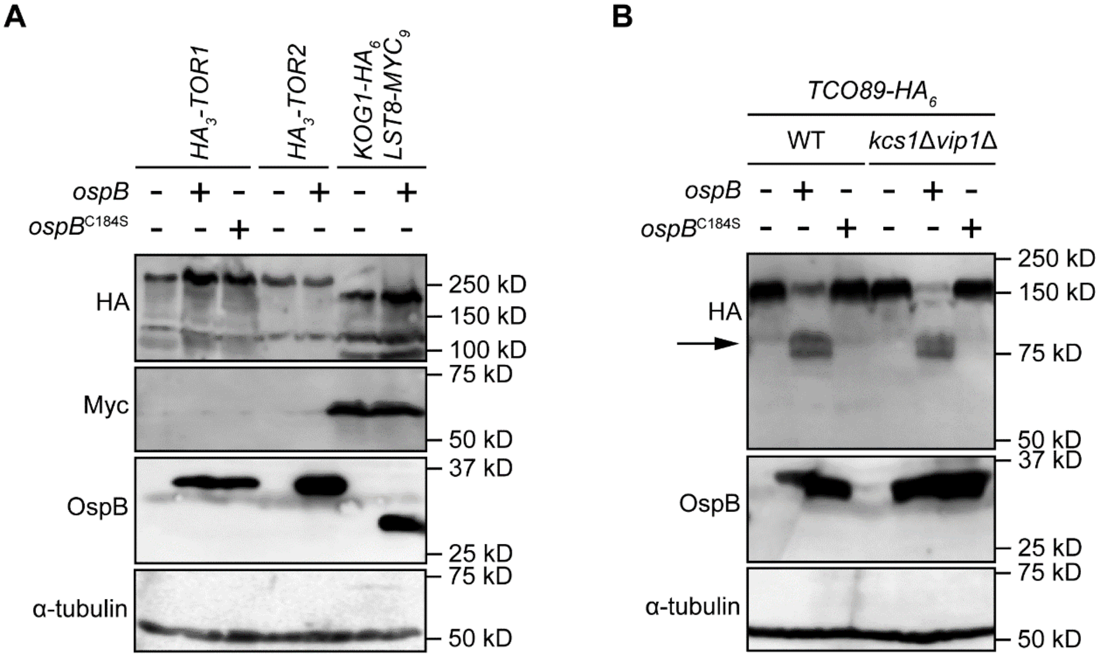


**FIGURE S7: Components of TORC1 other than Tco89p are not cleaved by OspB; inositol pyrophosphates are not required for Tco89p cleavage.**

(A) Western blot assessing cleavage of components of the yeast TORC1 complex by OspB. TORC1 proteins (Tor1p, Tor2p, Kog1p and Lst8) are tagged at the native loci. The OspB construct produced by the *KOG1-HA*$_6$ *LST8-myc*$_9$ strain is untagged, whereas the other *ospB*-epxressing strains produce C-terminally FLAG$_3$-His$_6$-tagged constructs. Alpha tubulin is the loading control (*n* =3).

(B) Western blot assessing cleavage of Tco89p in wild type and *kcs*1Δ*vip1*Δ yeast, in the presence of OspB, OspB(C184S) or vector control. Alpha tubulin is the loading control. The arrow marks the Tco89p C-terminal cleavage product (*n* = 3).

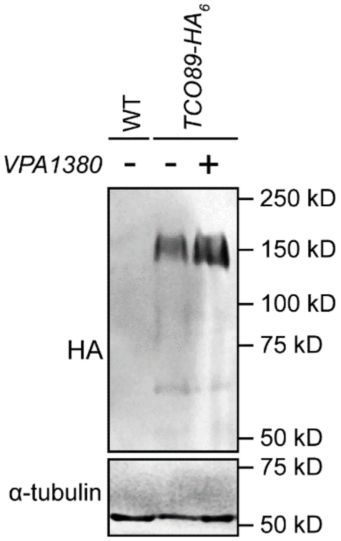


**FIGURE S8: VPA1380 does not cleave Tco89p.**

Western blot assessing cleavage of Tco89p by VPA1380. Alpha tubulin is the loading control (*n* = 2).

**TABLE S1: Yeast strains used in this study.**

| Strain | Description | Source |
|---|---|---|
| DH10B | Generic *E. coli* cloning strain | Laboratory collection |
| BY4741 | MATa *his3Δ1 leu2Δ0 met15Δ0 ura3Δ0* | Laboratory collection |
| *ipk1Δ* | MATa *his3Δ1 leu2Δ0 met15Δ0 ura3Δ0 ipk1::LEU2* | This study |
| *kcs1Δvip1Δ* | MATa *his3Δ1 leu2Δ0 met15Δ0 ura3Δ0 kcs1::kanMX4 vip1::LEU2* | This study |
| *TCO89-HA*$_6$ | *MATa his3Δ1 leu2Δ0 met15Δ0 ura3Δ0 TCO89-6xHA::hphNT1* | This study |
| *ipk1Δ TCO89-HA*$_6$ | MATa *his3Δ1 leu2Δ0 met15Δ0 ura3Δ0 ipk1::LEU2 TCO89-6xHA::hphNT1* | This study |
| *nta1ΔTCO89-HA*$_6$ | MATa *his3Δ1 leu2Δ0 met15Δ0 ura3Δ0 nta1::kanMX4 TCO89-6xHA::hphNT1* | This study |
| *gtr1Δgtr2Δ* | MATa *his3Δ1 leu2Δ0 met15Δ0 ura3Δ0 gtr1::kanMX4 gtr2::LEU2* | This study |
| *nta1Δ* | MATa *his3Δ1 leu2Δ0 met15Δ0 ura3Δ0 nta1::kanMX4* | Horizon MATa yeast knock-out collection |
| *ate1Δ* | MATa *his3Δ1 leu2Δ0 met15Δ0 ura3Δ0 ate1::kanMX4* | Horizon MATa yeast knock-out collection |
| *ubr1Δ* | MATa *his3Δ1 leu2Δ0 met15Δ0 ura3Δ0 ubr1::kanMX4* | Horizon MATa yeast knock-out collection |
| *rad6Δ* | MATa *his3Δ1 leu2Δ0 met15Δ0 ura3Δ0 rad6::kanMX4* | Horizon MATa yeast knock-out collection |
| *pib2Δ* | MATa *his3Δ1 leu2Δ0 met15Δ0 ura3Δ0 pib2::kanMX4* | Horizon MATa yeast knock-out collection |
| *tco89Δ* | MATa *his3Δ1 leu2Δ0 met15Δ0 ura3Δ0 tco89::kanMX4* | Horizon MATa yeast knock-out collection |
| *gid4Δ* | MATa *his3Δ1 leu2Δ0 met15Δ0 ura3Δ0 gid4::kanMX4* | Horizon MATa yeast knock-out collection |
| *gid10Δ* | MATa *his3Δ1 leu2Δ0 met15Δ0 ura3Δ0 gid10::kanMX4* | Horizon MATa yeast knock-out collection |
| *psh1Δ* | MATa *his3Δ1 leu2Δ0 met15Δ0 ura3Δ0 psh1::kanMX4* | Horizon MATa yeast knock-out collection |
| *ubc5Δ* | MATa *his3Δ1 leu2Δ0 met15Δ0 ura3Δ0 ubc5::kanMX4* | Horizon MATa yeast knock-out collection |
| *snf1Δ* | MATa *his3Δ1 leu2Δ0 met15Δ0 ura3Δ0 snf1::kanMX4* | Horizon MATa yeast knock-out collection |
| *gcn2Δ* | MATa *his3Δ1 leu2Δ0 met15Δ0 ura3Δ0 gcn2::kanMX4* | Horizon MATa yeast knock-out collection |
| *vam6Δ* | MATa *his3Δ1 leu2Δ0 met15Δ0 ura3Δ0 vam6::kanMX4* | Horizon MATa yeast knock-out collection |
| *KOG1-HA*$_6$ *LST8-MYC*$_9$ | MATa *his3Δ1 leu2Δ0 met15Δ0 ura3Δ0 ipk1::LEU2 KOG1-6xHA::hphNT1 LST8-9xMYC::natNT2* | (Chen *et al.*, 2020) |
| W303a *HA*$_3$*-TOR1* | MATa *leu2-3112 ura3-52 can1- 100 ade2-1 his3-11 trp1-11 HISMX6::3xHA-TOR1* | (Wedaman *et al.*, 2003) |
| W303a *HA*$_3$*-TOR2* | MATa *leu2-3112 ura3-52 can1- 100 ade2-1 his3-11 trp1-11 HISMX6::3xHA-TOR2* | (Wedaman *et al.*, 2003) |

**TABLE S2: Plasmids used in this study.**

| Plasmid | Purpose | Source |
|---|---|---|
| pAG413GAL-*ospB* | Yeast plasmid conditionally producing OspB | (Slagowski *et al.*, 2008) |
| pAG413GPD | Yeast centromeric parental vector | Susan Lindqvist |
| pAG413GPD-*EGFP* | Yeast plasmid producing EGFP | Susan Lindqvist |
| pAG413GPD-*ospB* | Yeast plasmid producing OspB | This study |
| pAG413GPD-*ospB*(C184S) | Yeast plasmid producing OspB(C184S) | This study |
| pAG413GPD-*ospB*(D108A) | Yeast plasmid producing OspB(D108A) | This study |
| pAG413GPD-*ospB*(D108A/D110A) | Yeast plasmid producing OspB(D108A/D110A) | This study |
| pAG413GPD-*ospB*(D109A) | Yeast plasmid producing OspB(D109A) | This study |
| pAG413GPD-*ospB*(D110A) | Yeast plasmid producing OspB(D110A) | This study |
| pAG413GPD-*ospB*(H144A) | Yeast plasmid producing OspB(H144A) | This study |
| pAG415 | Yeast centromeric parental vector | Susan Lindqvist |
| pAG415GAL-*ospB* | Yeast plasmid conditionally producing OspB | This study |
| pAG415-*NTA1* | Yeast plasmid expressing *NTA1* with native promoter and terminator | This study |
| pAG415-*nta1*(C187S) | Yeast plasmid expressing *nta1*(C184S) with native promoter and terminator | This study |
| pAG415-*nta1*(C187S)-*$HA_3$* | Yeast plasmid expressing *nta1*(C184S) from native promoter | This study |
| pAG415-*NTA1*-*$HA_3$* | Yeast plasmid expressing *NTA1* from native promoter | This study |
| pAG415-*RAD6* | Yeast plasmid expressing *RAD6* with native promoter and terminator | This study |
| pAG415-*rad6*(C88S) | Yeast plasmid expressing *rad6*(C88S) with native promoter and terminator | This study |
| pAG415-*TCO89*-*$HA_3$* | Yeast plasmid expressing *TCO89* from native promoter | This study |
| pAG415GAL-*VPA1380* | Yeast plasmid conditionally producing VPA1380 | This study |
| pCMV-*FLAG* | Mammalian expression vector | Laboratory collection |
| pCMV-*$FLAG_3$*-*TCO89* | Mammalian expression vector expressing *TCO89* | This study |
| pCMV-*myc* | Mammalian expression vector | Laboratory collection |
| pCMV-*myc*-*ospB* | Mammalian expression vector producing OspB | This study |
| pCMV-*myc*-*ospB*(C184S) | Mammalian expression vector producing OspB(C184S) | This study |
| pRS313 | Yeast centromeric parental vector | (Sikorski and Hieter, 1989) |
| pRS313-*IPK1* | Yeast plasmid expressing *IPK1* with native promoter and terminator | This study |
| pRS316 | Yeast centromeric parental vector | (Sikorski and Hieter, 1989) |
| pRS316GAL-*GST*-*$His_6$* | Yeast plasmid for conditional expression | This study |
| pRS316GAL-*GST*-*$His_6$*-*BRE1* | Yeast plasmid for conditional expression of *BRE1* | This study |
| pRS316GAL-*GST*-*$His_6$*-*bre1*(C663S) | Yeast plasmid for conditional expression of *bre1*(C663S) | This study |
| pRS316GAL-*ospB*-*$FLAG_3$*-*$His_6$* | Yeast plasmid conditionally producing OspB | This study |
| pRS316GAL-*VPA1380*-*$FLAG_3$*-*$His_6$* | Yeast plasmid conditionally producing VPA1380 | This study |
| pRS425 | Yeast 2-micron vector | (Christianson *et al.*, 1992) |
| pRS425GPD-*RFP*-*TCO89*-*$FLAG_3$* | Yeast multicopy vector overexpressing *TCO89* | This study |
| pYM16 | Yeast vector providing template for chromosomal insertion of *6xHA* tag | Euroscarf |

**TABLE S3: Primers used in this study.**

Restriction enzyme sites incorporated into primer sequences are underlined.

| Primer | Sequence | Purpose |
|---|---|---|
| HE107 | ACCAGTCGAAAATTGTCAGAGATAAGTTCCTTTTTTGAAAAGAAAGATCGTAACTGTGGGAATACTCAGG | Forward primer for amplification of *LEU2* marker from pRS315 to insert into *IPK1* locus |
| HE108 | TAATGTATGTGCATCTGCCAGTACCAAAGGTGGAAAGAAAAGTATACAGTTTTAAGCAAGGATTTTC | Reverse primer for amplification of *LEU2* marker from pRS315 to insert into IPK1 locus |
| IPK1_F | CACGTAGGAAAGCGA | Forward screening primer for generation of *ipk1* mutant |
| IPK1_R | CCCTTCGTTGAATATCG | Reverse screening primer for generation of *ipk1* mutant |
| HE088 | CATACAAATTCAAAAGCATCTCGTAGCATATTAATATATTGCAGAAGGTCGAGCAGACAAGCCCGTCAG | Forward primer for amplification of *LEU2* marker from pRS315 to insert into *VIP1* locus |
| HE090 | TAAATACTTATTTAGTTTTGGGTTACTAAATTAAAAATTGGGTGTGATCACTCCTTACGCATCTGTG | Reverse primer for amplification of *LEU2* marker from pRS315 to insert into *VIP1* locus |
| VIP1_F | TAGCAATCTCATCGCG | Forward screening primer for generation of *vip1* mutant |
| VIP1_R | CGAAAACTCCGGACCTAAA | Reverse screening primer for generation of *vip1* mutant |
| TW187 | ATGGAAAATGTGGGCTACATGCATACACAGCCACAACAAAGTCGTACGCTGCAGGTC | Forward primer for tagging endogenous *TCO89* gene at 3' -end |
| TW188 | ATTAGCTACTCTTTTAAACTGTGTGCTTCGTGTTGGTTGTTTGGAGACCGGCAGATCC | Reverse primer for tagging endogenous *TCO89* gene at 3' -end |
| TW217 | TAACAAAACTCCAGGACAACGGTACTAATACACATACAACTAACTGTGGGAATACTCAGG | Forward primer for amplification of *LEU2* marker from pRS315 to insert into *GTR2* locus |
| TW218 | TCTATATACCCTAATATTTTCATGCCTTACGTCTTCTTTTTTAAGCAAGGATTTTCTTAA | Reverse primer for amplification of *LEU2* marker from pRS315 to insert into *GTR2* locus |
| TW219 | AACCGATTAACATCCACAGA | Forward screening primer for generation of *gtr2* mutant |
| TW220 | AAAACTTTGGCACCTCTGTA | Reverse screening primer for generation of *gtr2* mutant |
| HE008 | GTAAAACGACGGCCAGT | Forward screening primer for yeast pRS suite vectors (M13F) |
| HE009 | GGAAACAGCTATGACCATG | Reverse screening primer for yeast pRS suite vectors (M13R-like) |
| HE001 | CCGTTTTACTTCAAGCGGCTCCGCTGATAAAGTGG | Forward primer for amplification of *ospB* (C184S) mutation by Quikchange |
| HE002 | CCACTTTATCAGCGGAGCCGCTTGAAGTAAAACGG | Reverse primer for amplification of *ospB* (C184S) mutation by Quikchange |
| HE003 | GGTTTATATTCTTGGGGCCGGTAGTCCTGGTTCTCATC | Forward primer for amplification of *ospB* (H144A) mutation by Quikchange |
| HE004 | GATGAGAACCAGGACTACCGGCCCCAAGAATATAAACC | Reverse primer for amplification of *ospB* (H144A) mutation by Quikchange |
| TW017 | TAGTAATAAATAATGCTGATGACGCAT | Forward primer for amplification of *ospB* (D108A) mutation by SOEing PCR |
| TW018 | ATGCGTCATCAGCATTATTTATTACTA | Reverse primer for amplification of *ospB* (D108A) mutation by SOEing PCR |
| TW311 | TAAATAATGATGCTGACGCATTGAA | Forward primer for amplification of *ospB* (D109A) mutation by SOEing PCR |
| TW312 | TTCAATGCGTCAGCATCATTATTTA | Reverse primer for amplification of *ospB* (D109A) mutation by SOEing PCR |
| TW313 | TAAATAATGATGATGCCGCATTGAA | Forward primer for amplification of *ospB* (D110A) mutation by SOEing PCR |

| | | |
|---|---|---|
| TW314 | TTCAATGCGGCATCATCATTATTTA | Reverse primer for amplification of *ospB* (D110A) mutation by SOEing PCR |
| TW315 | TAAATAATGCTGATGCCGCATTGAAT | Forward primer for amplification of *ospB* (D108A/D110A) double mutation by SOEing PCR |
| TW316 | ATTCAATGCGGCATCAGCATTATTTA | Reverse primer for amplification of *ospB* (D108A/D110A) double mutation by SOEing PCR |
| TW001 | TAGCGCCGTCTTTCAGTTCG | Forward screening primer for *nta1* mutant |
| HE104 | AAACATCTACAACATTGCTTCACAA | Reverse screening primer for *nta1* mutant |
| TW002 | CACCAGCTTCGTCGTCCATC | Forward screening primer for *ate1* mutant |
| TW003 | TCGTTTTCACCCCGCGTATT | Reverse screening primer for *ate1* mutant |
| TW004 | ATCATCGTCGTCTCCATCGC | Forward screening primer for *ubr1* mutant |
| TW005 | CCCAGGCGCTACTAAGACCA | Reverse screening primer for *ubr1* mutant |
| TW006 | GCACACGTCGCTAGAACCAA | Forward screening primer for *rad6* mutant |
| TW007 | TGCCCGCACAGAAGAGTACT | Reverse screening primer for *rad6* mutant |
| TW087 | ATG<u>CTGCAGGAATTCGAGCTC</u>TAGCGCCGTCTTTCAGT | Forward primer to amplify *NTA1* with native promoter |
| TW088 | TCA<u>CTCGAG</u>TCA<u>GGATCC</u>CCTAAACACTTCAAATTGGACC | Reverse primer to amplify *NTA1* without stop codon for tagging |
| TW089 | AAGTCCATAGAAATACCTATTGATG | Forward primer for amplification of *nta1* (C187S) mutation by SOEing PCR |
| TW090 | GCATCAATAGGTATTTCTATGGAC | Reverse primer for amplification of *nta1* (C187S) mutation by SOEing PCR |
| TW208 | TCA<u>CTCGAG</u>TCA<u>GGATCC</u>CGTTCCCTTATCCTTCGG | Reverse primer for amplification of *NTA1* with native terminator and stop codon |
| MY009 | ATG<u>GAGCTC</u>GGATATGGTACCGATGTTGT | Forward primer to amplify *RAD6* with native promoter |
| MY010 | TCA<u>CTCGAGAAGCTT</u>CTATCA<u>TGATCA</u>GTCTGCTTCGTCGT | Reverse primer for amplification of *RAD6* with stop codon |
| MY001 | GCAAATGGTGAAATTTCTTTGGATAT | Forward primer for amplification of *rad6* (C88S) mutation by SOEing PCR |
| MY002 | TGCAAAATATCCAAAGAAATTTCACC | Reverse primer for amplification of *rad6* (C88S) mutation by SOEing PCR |
| MY007 | ATG<u>ATCGAT</u>AGATATGACGGCCGAGC | Forward primer for amplification of *BRE1* |
| MY008 | TCA<u>GGGCCC</u>TTACAAGTGCACTGTCAATAAATC | Reverse primer for amplification of *BRE1* with stop codon |
| MY003 | CTATTAAAACCTCTGGCCATGT | Forward primer for amplification of *bre1* (C663S) mutation by SOEing PCR |
| MY004 | AGACATGGCCAGAGGTTT | Reverse primer for amplification of *bre1* (C663S) mutation by SOEing PCR |
| TW113 | ATG<u>GAGCTC</u>GGCTGGCACGATGATTAA | Forward primer to amplify *TCO89* with native promoter |
| TW114 | TCA<u>GCGGCCGC</u>C<u>ACTAGT</u>CCTTTGTTGTGGCTGTGT | Reverse primer to amplify *TCO89* without stop codon for tagging |
| HE041 | TGTAGTCGATGTCATGATCCTTGTAATCACCGTCATGGTCCTTGTAGTC<u>GCTAGC</u>ATCCAGTTCTTTATTAATAA | Reverse primer 1 for constructing *ospB-3xFLAG-6xHis* |
| HE042 | TAGAGCGAT<u>AAGCTT</u>TCAACCATGGTGATGGTGATGATGCTTGTCATCGTCATCCTTGTAGTCGATGTCATGATC | Reverse primer 2 for constructing *ospB-3xFLAG-6xHis* |
| TW222 | ATG<u>TGTACA</u>AG<u>GATATCAGGCCT</u>GTTCATCGAGGAAGGACTTT | Forward primer to amplify *TCO89* without start codon |
| TW223 | TCA<u>GTTAACGCGGCCGCGCTAGCTCTAGA</u>TCACCTTTGTTGTGGCTG | Reverse primer to amplify *TCO89* with stop codon for tagging |

| | | |
|---|---|---|
| TW189 | ATGGAGCTCCCCCACCTCACAAATCTATT | Forward primer for amplification of *IPK1* with native promoter |
| TW190 | TCACTCGAGTCCCTTACATCCCAATCTTTG | Reverse primer for amplification of *IPK1* with stop codon and native terminator |
| TW232 | ATGTCTAGAGTCGACTCCGGTTCTGCTGCTAGTGGTATGG CCTCCTCCGAG | Forward primer for amplification of *RFP* for *TCO89* overexpression |
| TW233 | TCAGGTACCATCGATAGATCTTCACTACTTGTCATC GTCATC | Reverse primer for amplification of 3x*FLAG* tag for *TCO89* overexpression |
| KW003 | TCAGCTAGCACTAGTATCTAAATCAGATTCTAAGGT AAC | Reverse primer for amplification of *VPA1380* without stop codon |

**TABLE S4: Genes identified as required for OspB-dependent sensitization of yeast to caffeine.**

Genes encoding proteins with a characterized role in TORC1 signaling are shown in bold.

| Gene | Locus Tag | Gene | Locus Tag | Gene | Locus Tag | Gene | Locus Tag |
|---|---|---|---|---|---|---|---|
| - | *YBR085C-A* | *CIT3* | *YPR001W* | *MFA2* | *YNL145W* | *SAK1* | *YER129W* |
| - | *YDR442W* | *CTF19* | *YPL018W* | *NAT3* | *YPR131C* | ***SAP185*** | ***YJL098W*** |
| - | *YEL025C* | *DCV1* | *YFR012W* | *NGR1* | *YBR212W* | *SAP30* | *YMR263W* |
| - | *YGL041C-B* | *DOA1* | *YKL213C* | ***NPR1*** | ***YNL183C*** | *SIF2* | *YBR103W* |
| - | *YGR025W* | *DSK2* | *YMR276W* | *NTA1* | *YJR062C* | *SKI8* | *YGL213C* |
| - | *YLR434C* | *ECM23* | *YPL021W* | *PAF1* | *YBR279W* | *SNA3* | *YJL151C* |
| - | *YMR052C-A* | *ETR1* | *YBR026C* | *PBI2* | *YNL015W* | *SOD1* | *YJR104C* |
| - | *YNL040W* | *FOX2* | *YKR009C* | *PDB1* | *YBR221C* | *SOL1* | *YNR034W* |
| - | *YNL057W* | ***FPR1*** | ***YNL135C*** | *PEP12* | *YOR036W* | *SPO7* | *YAL009W* |
| - | *YNL140C* | ***GLN3*** | ***YER040W*** | *PEP7* | *YDR323C* | ***STP1*** | ***YDR463W*** |
| - | *YNL195C* | *HAT1* | *YPL001W* | *PET191* | *YJR034W* | *SWA2* | *YDR320C* |
| - | *YNL296W* | *HOS2* | *YGL194C* | *PUB1* | *YNL016W* | *SYF2* | *YGR129W* |
| - | *YNR005C* | *HSE1* | *YHL002W* | *RAD6* | *YGL058W* | ***TIP41*** | ***YPR040W*** |
| *AIM22* | *YJL046W* | *HTD2* | *YHR067W* | *RBS1* | *YDL189W* | *UBA3* | *YPR066W* |
| *ALF1* | *YNL148C* | *IPK1* | *YDR315C* | *RIB1* | *YBL033C* | *UBP3* | *YER151C* |
| *APE2* | *YKL157W* | *IRC21* | *YMR073C* | *RNR1* | *YER070W* | *UBR1* | *YGR184C* |
| *ATE1* | *YGL017W* | *IWR1* | *YDL115C* | *RPN4* | *YDL020C* | *VIP1* | *YLR410W* |
| *BRE5* | *YNR051C* | *KCC4* | *YCL024W* | *RPS12* | *YOR369C* | *VPS13* | *YLL040C* |
| *BRR1* | *YPR057W* | *LIP2* | *YLR239C* | ***RRD1*** | ***YIL153W*** | | |
| ***BUL1*** | ***YMR275C*** | *LPD1* | *YFL018C* | ***RTG3*** | ***YBL103C*** | | |
| *CBT1* | *YKL208W* | *MCT1* | *YOR221C* | *SAC3* | *YDR159W* | | |

**TABLE S5: Genes identified as suppressors of OspB-dependent sensitization of yeast to caffeine.**

| Gene | Locus Tag | Gene | Locus Tag | Gene | Locus Tag |
|---|---|---|---|---|---|
| - | *YAR023C* | *FMT1* | *YBL013W* | *QCR6* | *YFR033C* |
| - | *YBL100C* | *GAL11* | *YOL051W* | *RIM11* | *YMR139W* |
| - | *YBR116C* | *HIS3* | *YOR202W* | *SEC3* | *YER008C* |
| - | *YBR284W* | *HSP60* | *YLR259C* | *SIW14* | *YNL032W* |
| *BRE1* | *YDL074C* | *JJJ3* | *YJR097W* | *SLX8* | *YER116C* |
| *CLN3* | *YAL040C* | *LAT1* | *YNL071W* | *TCB1* | *YOR086C* |
| *COS3* | *YML132W* | *LSB1* | *YGR136W* | *TKL2* | *YBR117C* |
| *CSM1* | *YCR086W* | *MTC6* | *YHR151C* | *UBC1* | *YDR177W* |
| *CYC3* | *YAL039C* | *PBY1* | *YBR094W* | *UMP1* | *YBR173C* |
| *DDI1* | *YER143W* | *PPE1* | *YHR075C* | *YUH1* | *YJR099W* |
| *DDP1* | *YOR163W* | *PRX1* | *YBL064C* | | |
| *EMA17* | *YIL029C* | *PTC3* | *YBL056W* | | |
